# Chioso: Segmentation-free Annotation of Spatial Transcriptomics Data at Sub-cellular Resolution via Adversarial Learning

**DOI:** 10.1101/2024.06.03.597195

**Authors:** Ji Yu

**Affiliations:** UConn Health, 400 Farmington Ave, Farmington, CT, 06030

**Keywords:** Cadmium, Connexin, Rictor, mTOR2, Oxidative stress, Autophagy

## Abstract

Recent advances in spatial transcriptomics technology have produced full-transcriptomic scale dataset with subcellular spatial resolutions. Here we present a new computational algorithm, chioso, that can transfer cell-level labels from a reference dataset (typically a single-cell RNA sequencing dataset) to a target spatial dataset by assigning a label to every spatial location at sub-cellular resolution. Importantly, we do this without requiring single cell segmentation inputs, thereby simplifying the experiments, and allowing for a more streamlined, and potentially more accurate, analysis pipeline. Using a generative neural network as the underlying algorithmic engine, chioso is very fast and scales well to large datasets. We validated the performance of chioso using synthetic data and further demonstrated its scalability by analyzing the complete MOSTA dataset acquired using the Stereo-Seq technology.

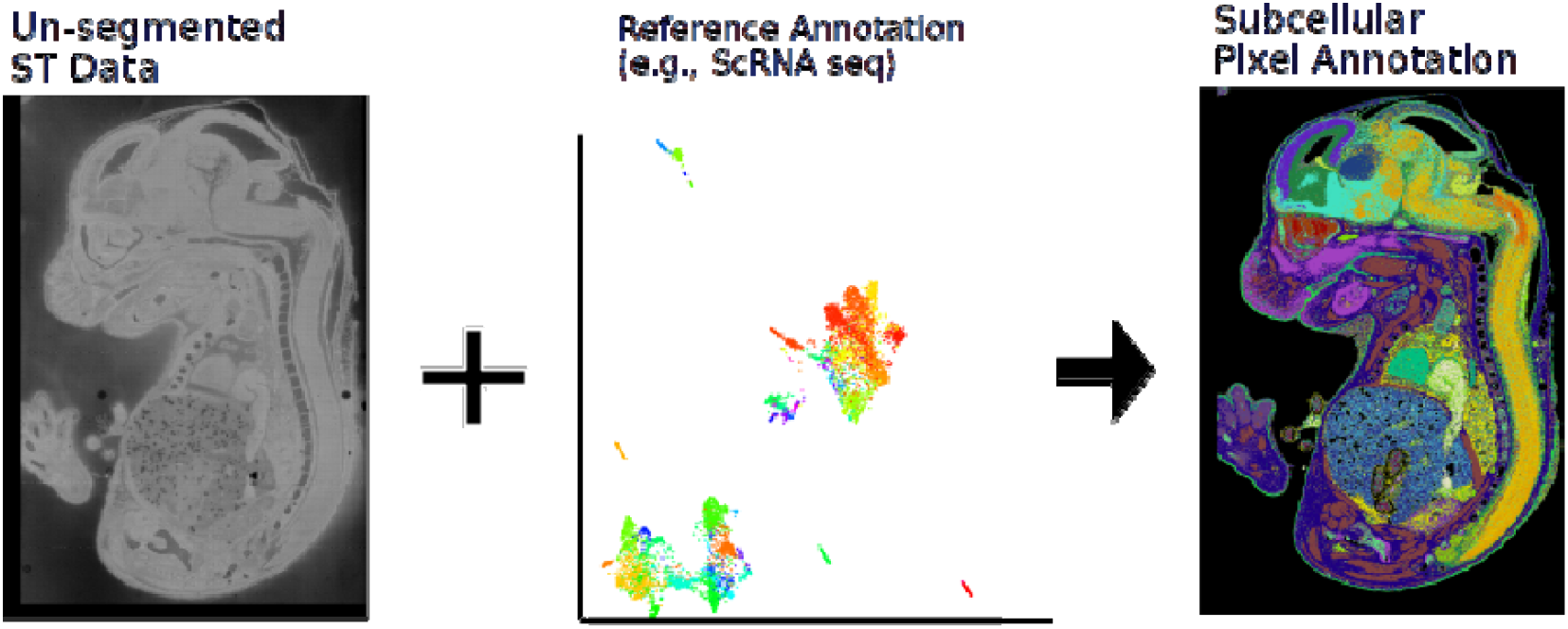

## Main

Spatially resolved transcriptomics (ST)^1,2^ provide high dimensional view of cells’ transcriptional program and cell states in a spatially resolved manner. These technologies are powerful in elucidating the complex tissue structures and reveals spatial organization of specific cellular subtypes. To this end, a very useful analysis is the reference-based annotation, which transfers existing biological knowledge, such as cell types or cell states, built upon a reference dataset, i.e., single-cell RNA sequence (scRNASeq), to the ST dataset. The assay thereby lend data interpretability to the otherwise opaque ST data^3–5^. Such analysis further enables comparisons of multiple experiments, which can potentially be using different types of ST technologies.

The major challenge in reference-based labeling, from a computational biology perspective, arises from the mismatched supports of the ST dataset and the reference dataset: The reference dataset are typically measurements of individual cells, whereas the ST datasets are measurements of locations, which does not match to a cell in a 1:1 fashion. There has been a significant amount of work published already dealing with the case where each spatial location corresponds to a few cells^6–15^. However, in recent years, we have seen a concerted push towards acquiring ST data at true microscopic, sub-cellular spatial resolutions^16–22^, in which case, each measured location contains only a small fraction of the total RNAs expressed by the cell. Standard procedure in this case is to first segment the ST data based on a secondary image of the cells, which added significant amount of additional experimental and computational complexities^23–27^ to the pipeline. In addition, for capture-based ST technologies, it is possible for the detected RNA locations to be outside the physical cell boundaries, due to the diffusion process during capture. In such scenario, even a perfect segmentation may produce imperfect quantifications of gene expressions.

Recognizing the problems of the segmentation approach, some authors have previously suggested alternative, segmentation-free approaches: for example, Park et al.^28^ proposed to compute spatially smoothened expression profiles using kernel density estimation and use local maxima to approximate cell expressions. Mages et al.^8^ proposed to perform label assignment for every RNA detection by computing integrated expressions over multiple spatial filters and selecting the final answer based on a voting algorithm. However, none of the proposed methods can scale to full transcriptome-scale ST data, and therefore are limited to smaller datasets.

Here we propose a new algorithm to perform reference-based labeling by assigning a label to every pixel/spatial location in the ST data, where the pixel size is significantly smaller than that of a cell. We show that the accurate labeling can be obtained without needing single cell segmentation input. The underlying engine for the algorithm is a neural network (NN) trained using adversarial learning. To demonstrate the scalability of the algorithm, we performed labeling experiments on the MOSTA^16^ dataset, a large ST dataset of developing whole mouse embryo with ∼20 billion total RNA reads, acquired using the StereoSeq technology. We labeled the complete dataset at a 2-µm pixel-size (∼ 0.9 billion total pixels labeled) in under 6 hours (combined training and inference time) on a single GPU.

## Results

### Algorithm outline

Fig. 1 shows the high-level overview of the chioso algorithm and its top-level components (Table S1). The overall computational workload can be divided into three sub-steps:

**Figure 1.**
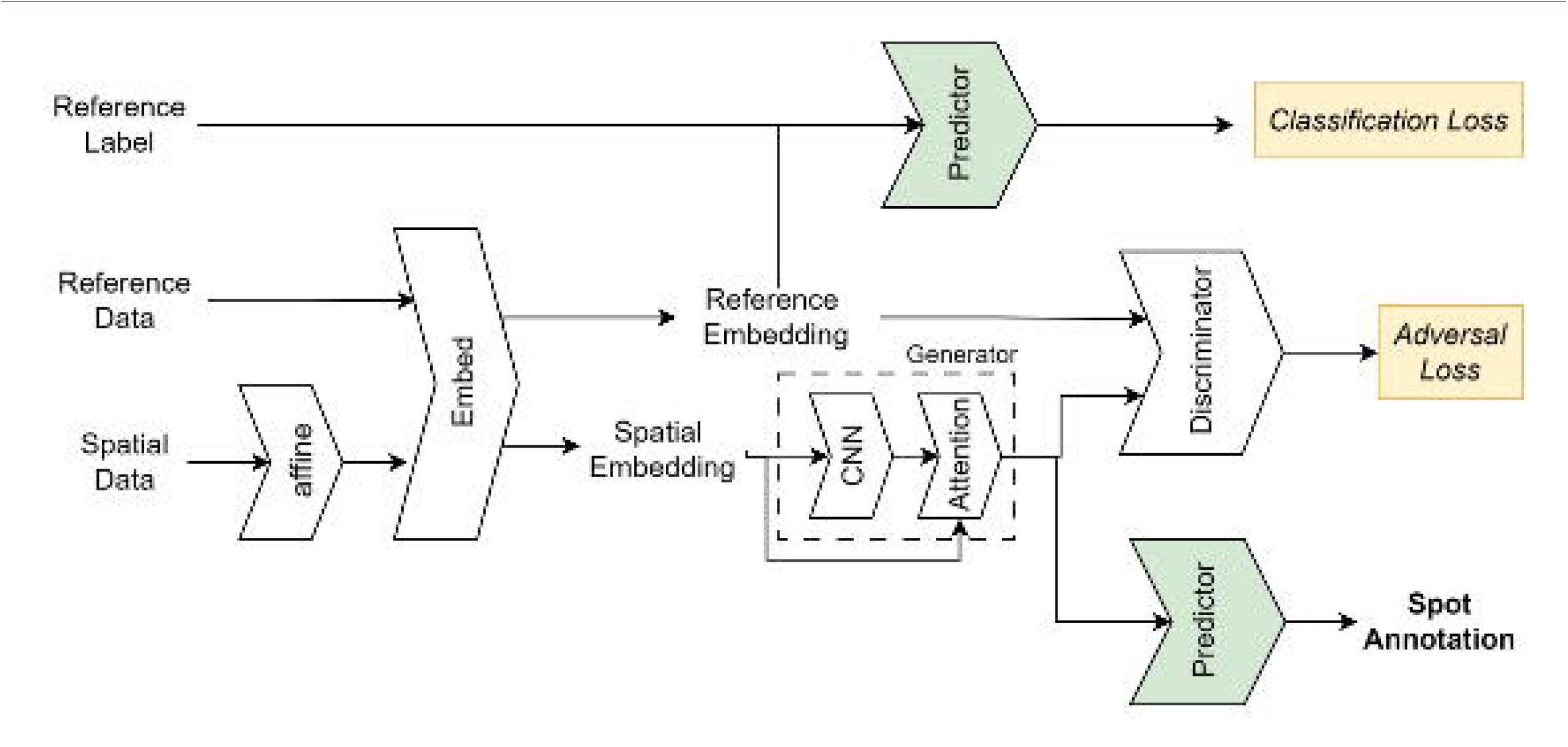
Outline of the chioso algorithm showing its sub-components. Chioso is a deep-learning based algorithm using the adversarial learning training scheme. Model parameters are obtained by co-training two competing modules: the generator, which synthesize whole-cell expression profile based on noisy pixel-level expression data, and the discriminator, which try to distinguish synthesized expression profiles from real experimental single-cell expression profiles.

1. Predictor training using the reference dataset. The purpose of this step is to train a neural network classifier to predict cell labels from the scRNAseq inputs. For computational efficiency, a simple linear embedding module (Fig. 1) was employed to reduce the dimension of the input data (i.e., number of genes, >20,000) to a manageable scale (several hundreds). This is then followed by a nonlinear classifier (predictor in Fig.1), implemented as a multi-layer perceptron (MLP) to output probability of each label.
2. Generator training using the combined reference and ST datasets. The key idea of the chioso algorithm is to learn a NN function, i.e., a “generator”, that can synthesize, at each pixel location, a plausible whole-cell expression profile. Note that the generator shares the same embedding module as that of the predictor. We design the generator as a simple attention mechanism that integrate RNA reads from a nearby neighborhood of pixels. However, instead of using a pre-defined spatial filter for integration, chioso tries to learn the attention weights directly from the data via a convolutional neural network (CNN, Fig. 1). To train such a model, we use the adversarial training scheme, in which the generator is co-trained with an additional neural network, i.e., a discriminator (Fig. 1). The discriminator will learn to distinguish the “synthesized” expression profile from a “real” one, sampled from the reference dataset; while at the meantime, the generator will learn to get better at the synthesis job to “fool” the discriminator. The end-result is that the generator will learn the correct weights corresponding how likely is a neighboring pixel belonging to the same cell.
3. Inference. Finally, the trained generator can be applied to all ST data to predict at each pixel what is the likely whole-cell expression profile. Once those results were obtained, it is straightforward to employ the predictor trained in the first step to generator cell-type labels for each pixel.

In addition, the model also tries to account for domain drifts due to the platform effect, such as those described by Cable et al.^6^, by employing an affine transformation (Fig. 1) to the ST input. These platform effects include differences in read depth between the reference technology and the targeting ST technology, and different bias in library preparation and capture rates of individual genes, etc. However, unlike previous algorithm, chioso does not pre-computing these factors relying on a specific statistical model, but instead try to learn the parameters from the input data as part of the adversarial training step.

Finally, because the model output is probabilistic, we provided an optional Markov random field module to encourage short range consistency the final labeling output. Similar techniques have been previously shown to increase labeling accuracy in cell level ST data analysis^29^. Here we implement the module using a fast dense conditional random field implementation by Krahenbuhl et al^30^.

### Validation with synthetic ST data

To validate chioso algorithm and evaluate its performance, we generated synthetic ST data with known ground truth labeling based on computer simulation. We first took a previously produced^31^ single-cell segmentation dataset and extracted all segmentation masks (500 segmented images). The dataset was produced by segmenting experimentally acquired optical microscopic images of A431 cells. Next, we randomly sampled from a reference scRNAseq dataset (TOME)^32^ expression profiles of three different cell types (i.e., spinal cord excitatory neuron, inhibitory neuron, and motor neuron), and distributed the RNAs into individual cells in the segmented images. The three cell types were sampled at a ratio of 15:10:1 to create imbalances in the cell populations. Finally, to mimic RNA location error in ST experiments, we let each RNA molecule to randomly diffuse with a mean square displacement of 72 µm^2^, potentially move crossing the cell boundaries. An example of the resulting synthetic data is shown in Fig. 2a.

**Figure 2.**
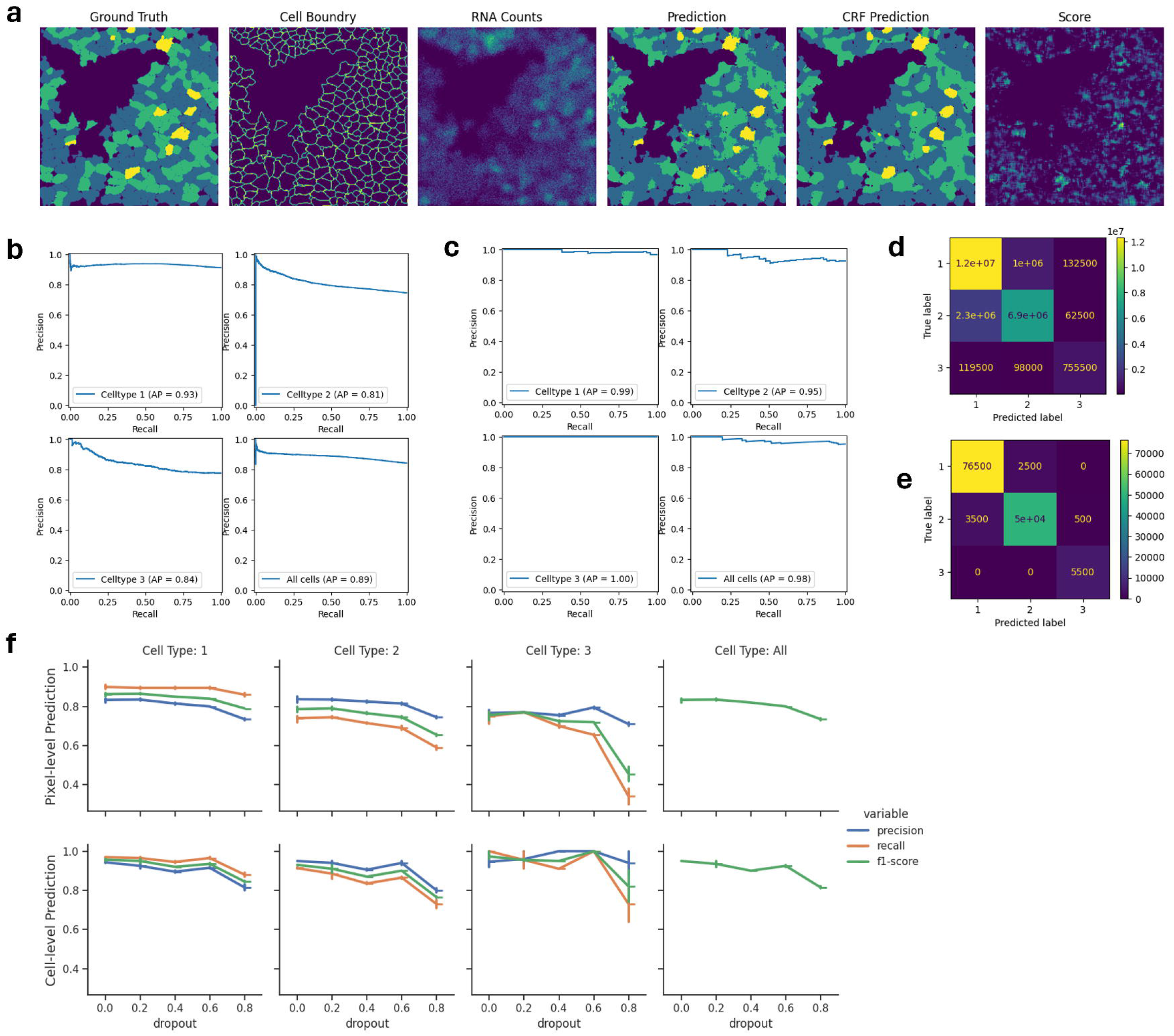
Validation of chioso using synthetic ST data. (**a**) An example of the synthetic data and corresponding chioso prediction. From left to right, the images shown represent ground truth labeling of three cell types, the boundaries of individual cells, the raw RNA counts of the synthetic ST data, the chioso predicted pixel label, the chioso prediction in combination with a CRF model, and discriminator loss value of each pixel, which we use to score the predictions. Scale bar corresponds to 20 µm. (**b**) Precision-recall curves of the chioso prediction on the whole dataset (500 images), ranked by the prediction scores. All curves exhibited decaying patterns, consistent with the expectation that the scores are positively correlated with the prediction accuracy. Average Precision scores (area under the curve) are shown for each cell type as well as the aggregate. (**c**) Same as **b**, but for cell-level label predictions, which were obtained via a voting mechanism. See text for details. (**d**) Confusion matrix for the pixel-level predictions. (**e**) Confusion matrix for the cell-level predictions. (**f**) Evaluation metrics of chioso predictions on dropout datasets, in which fractions of RNAs were randomly removed from the input data. The plots show the precision, recall and F1 scores (the geometric mean of the precision and the recall) for all three cell types as well as the aggregated F1 score for all cell types, Top row is the results for pixel-level predictions and the bottom row is for cell-level predictions.

**Figure 3.**
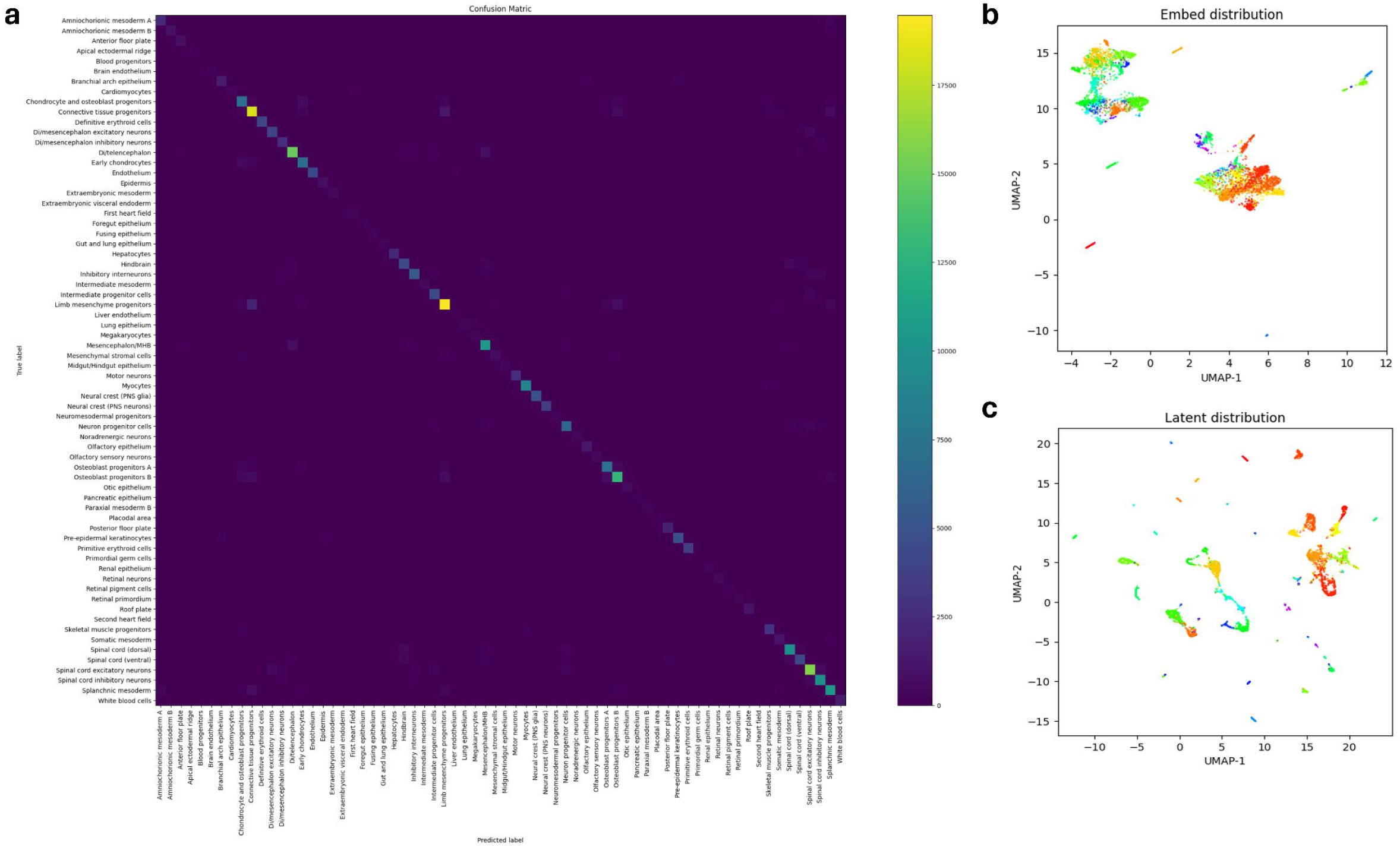
Evaluation of the neural-network based cell-type classifier using TOME dataset. (**a**) Confusion matrix of the classifier using the validation split (20%) of the input dataset. The classifier takes single-cell RNA expression profile as input and classify the cell into one of the 68 potential cell types (labels in a). (**b**) The UMAP visualization of the distribution of the input dataset. (**c**) The UMAP visualization of the latent representations of the single cell data after nonlinear projection by the neural network. The data shows effective separation of different clusters in the project space.

We then ran the chioso pipeline to predict pixel labels at 1.6 µm resolution (Fig. 2). We found that prediction correctly captures the domain shapes and reproduced the spatial characteristics of the ground truth distribution (Fig 2a). We also explored whether we could use the discriminator loss as a scoring measure to rank the predictions from high to low confidence (Fig 2a). A quick visual examination indeed showed that the scores obtained are lower in image regions where there is no cell present, and higher in the cellular region. Furthermore, sorting the pixel labels based on the score values and plotting the precision vs recall curve (Fig 2b) showed decaying functions for all three cell types, as well as the aggregated results, indicating indeed that pixels with high scores have higher average prediction accuracy. These results confirmed that not only can chioso make label predictions at pixel level, but it also provided a probabilistic measure of the prediction confidence, potentially allowing users to reject low confidence predictions. Quantification of the pixel-level predictions (Fig. 2b) showed that the average precisions (APs) are 0.94, 0.81, and 0.84 for the three cell types, with higher AP for the most abundant cell type, and 0.89 for all three cell types aggregated.

To further evaluate chioso’s error characteristics, we also generated “cell-level” predictions by incorporating the known cell segmentation masks. For each cell, we examined all pixel-level labels within the cell area and produced one “cell-level” prediction by choosing the most frequent label. We then quantified the accuracies of the cell-level predictions (Fig. 2c) by computing the precision-recall curves, ranking the results by the average scores of each cell. As expected, cell-level predictions are significantly better than pixel-level results, with an aggregated AP metric of 0.98 for all cell types. To establish an upper bound of the cell-level prediction accuracy, we replace chioso’s generator module with a derteministic function that simply integrate pixel values based on the known segmentation masks. This modified “chioso” model, therefore, has the perfect knowledge of cell masks, but still try to learn the platform effects and use the same non-linear predictor as the verbatim chioso model. We found that the modified chioso model also achieves and AP score of 0.98. Therefore, it is likely that chioso cell-level predictions have reached or close to its theoretical limit.

### Robustness of predictions

One of the challenges in labeling ST data is that they are usually noisier than bulk or single-cell sequencing data. StereoSeq^16,33^, an emerging ST technology with sub-micrometer spatial resolution, for example, produce results with much lower read depth than scRNAseq. Therefore, we set out to evaluate how resistant is chioso against dropout noise. To do that, we randomly removed a fraction of RNAs, starting from 20% and up to 80%, from the synthetic data we generated, and re-ran the chioso prediction pipeline on the altered datasets.

As shown in Fig. 2f, we found that the chioso algorithm is resistant to dropout noise. At both the pixel level and cell level, the precision, recall and F1 score showed only minor drop for altered input data with up to 60% dropout. At 80% dropout, we saw a significant decrease in the recall score of the least abundant cell type (cell type III), although the precision scores remained relativity stable. The impacts on the two abundant cell types, even at 80% dropout, are significantly less pronounced.

### Evaluating chioso on MOSTA/TOME datasets

Next, we evaluated chioso on a realistic large scale ST dataset, namely, the MOSTA dataset^16^. The dataset contains Stereo-Seq results on 53 sections of whole mouse embryo, ranging from developmental stage E9.5 to E16.5, with ∼20 billion total RNA reads. While the original data was obtained using DNA balls at 0.5 µm spacing, we had performed labeling on a 2-µm grid by first binning the input data. For reference dataset, we used the previously published TOME dataset^32^, which encompasses experimental scRNAseq data from multiple experiments, covering mouse embryos from E3.5 to E13.5 developmental stages. The original authors have labeled ∼1.5 million cells using a set of 68 different cell type labels.

We first trained the predictor model using the 80% of the TOME dataset and validated the results using the remaining 20%. Because the population of different cell types are highly imbalanced, we evaluate the model using the balanced accuracy metric to avoid overemphasizing a few highly represented cell types. We achieved 0.88 balanced accuracy. We found that it is important to learn the embedding vectors directly from the data, as using a rationally designed dimension reduction scheme, such as one based on principal component analysis, resulted in decreased performance (Table S2). Another factor is the hidden dimension of the embedded representation. Higher dimensions typically increase the predictor capacity at the cost of higher computational time. We found that for TOME dataset the model performance increases when we increase the hidden dimension up to ∼256, and plateaus after that (Fig. S1). Finally, there are significant recent interests in modeling single-cell expression using Transformer type of model architecture^34–36^, which are significantly more expressive than MLP, but also much slower. We trained a transformer model on the same training data for comparison. We found that the accuracy of the transformer model is essentially the same (Table S2). Therefore, we concluded that a simpler MLP predictor is sufficient for our purpose. Training a MLP model is extremely fast. For the TOME dataset, the model can be trained in < 40 min.

Next, we trained the full chioso model using the adversarial training scheme. Because the reference dataset covers only embryo up to E13.5, during training we similarly sample only E9.5-E13.5 portion of the ST dataset, with a total training time of ∼4 hours. However, during inference, we employed the trained model to make prediction for all MOSTA data, i.e., E9.5 – E16.5, relying on the generalizability of the trained model. The inference step took about 1 hour, making the total time for the combined computational pipeline to be < 6 hours.

Examples of the prediction results are shown in Fig. 5. Embryos from E9.5-E13.5, i.e., data used during training, were shown in Fig 5a, whereas the rest are shown in Fig 5b. Pseudo-color visualization is compact but can be difficult to read when there are many different cell types, thus, we also visualize individual cell types as binary images (Fig. S2a-c). Overall, the chioso predictions are consistent with our existing knowledges of the embryo (Fig.4a). For example, cell types such as myocytes show large domain structures (Fig. 4a), consistent with the expectation that the corresponding tissue type (muscle) are comprised of mainly one cell type. On the other hand, in spinal cord, multiple cell types (e.g., different neuronal subtypes) can be found in an intermingled pattern, with domain sizes comparable to the sizes of single cells (Fig. 5a). We also found chioso capable to resolving small structures at distance scale corresponding to single cells. For example, it can resolve the monolayer of epidermis structure from the layer of keratinocyte precursor right underneath (Fig 5b). These qualitative evidence lent confidence to the validity of the chioso predictions.

**Figure 4.**
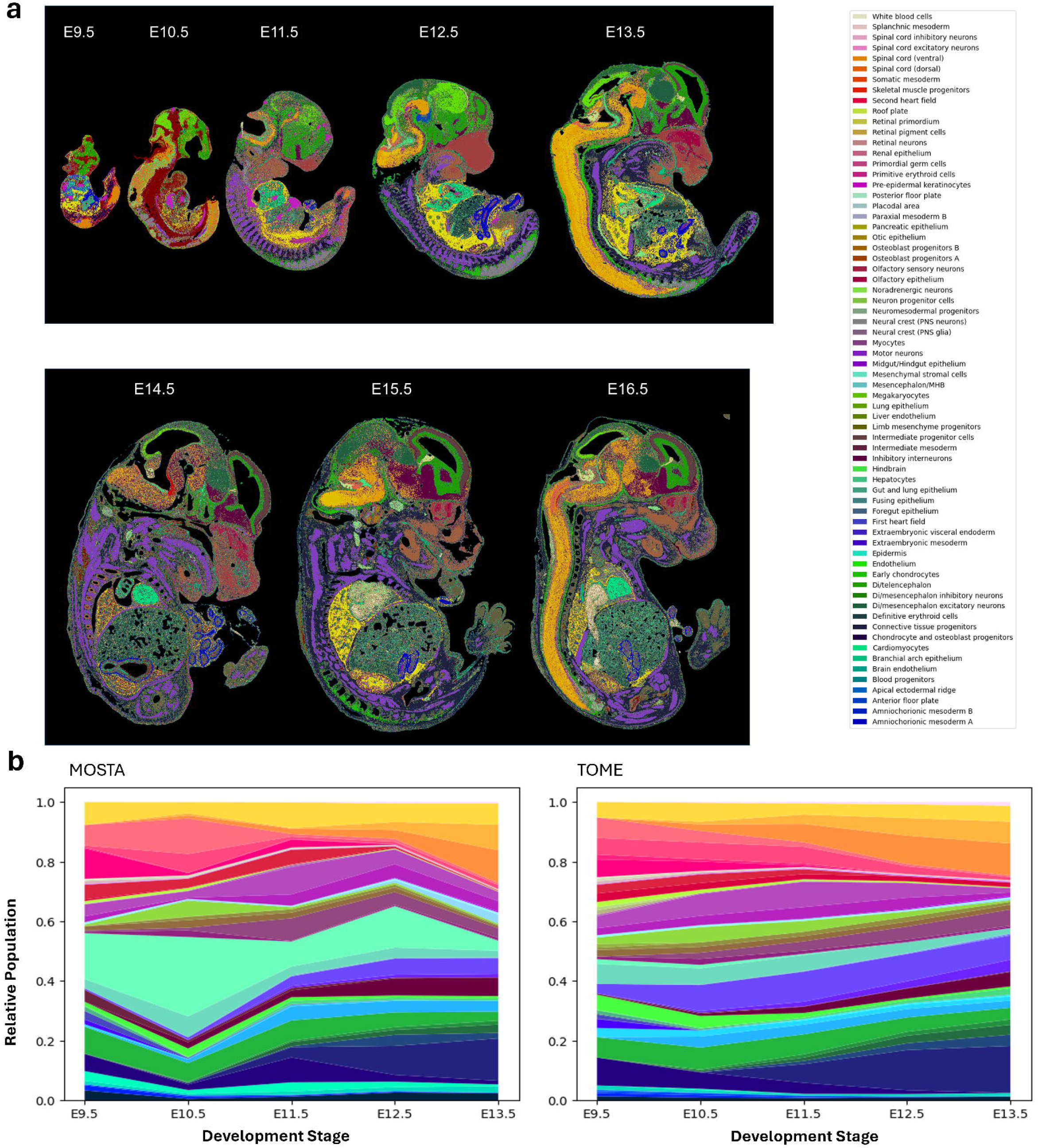
Chioso transfer cell type label to high-definition ST data without segmentation. (**a**) Examples of chioso labeling for MOSTA stereo-seq data using TOME dataset as the reference. The top row represents samples (E9.5-E13.5) matching the reference and used during model training. The bottom row (E14.5-E16.5) demonstrates the model generalization to mismatched input data. (**b**) Comparison of the relative cell populations of the MOSTA samples (mouse embryo sections), as inferred by the chioso labeling, and that of the TOME samples (whole mouse embryo).

**Figure 5.**
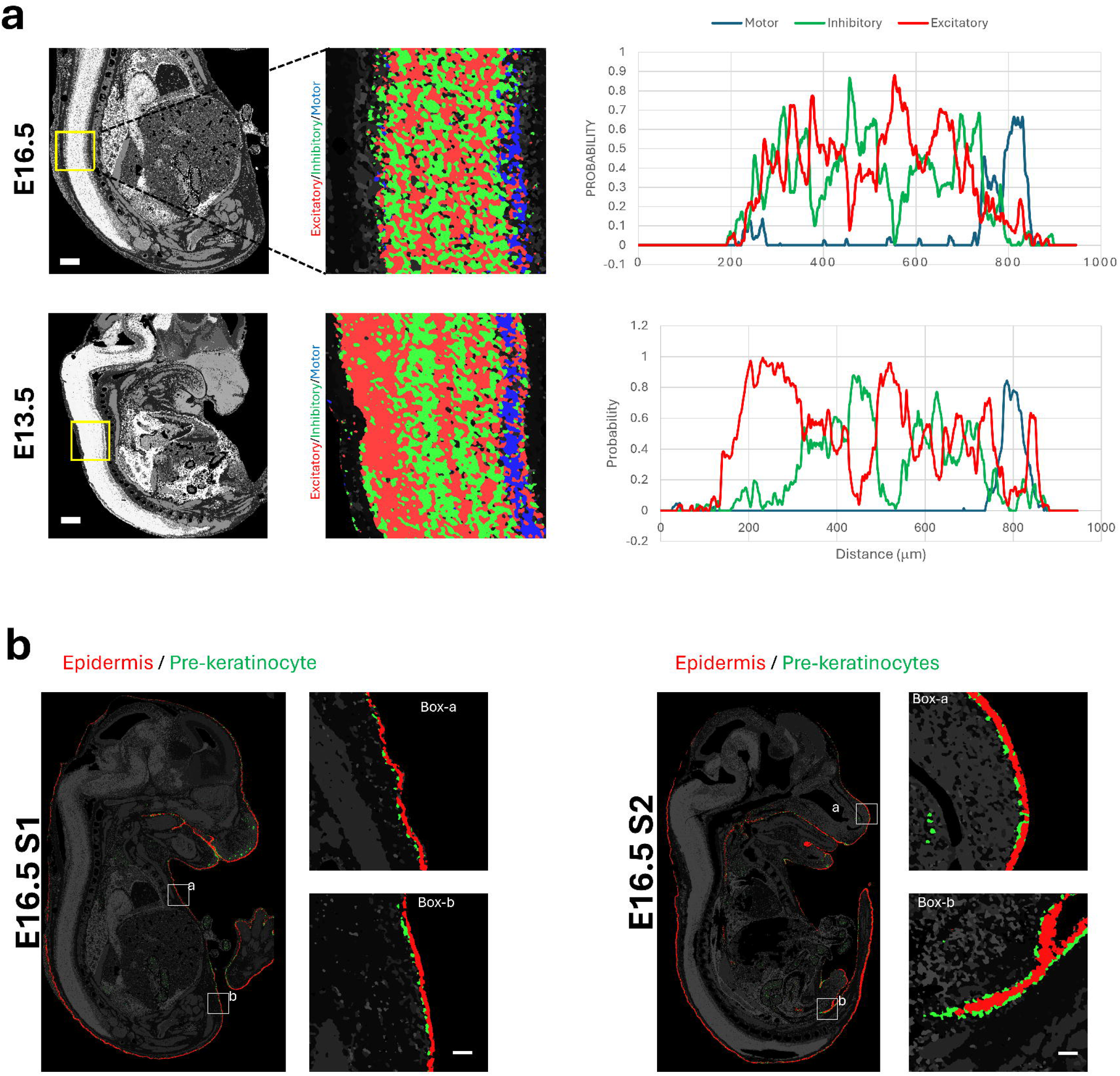
Chioso labeling reveals complex spatial organization of cells. (**a**) Chioso labeling of a E16.5 sample (top) and E13.5 sample (bottom) revealed the spatial organizations of three dominant cell types (excitatory neuron, inhibitory neuron and motor neuron) within the region. Zoomed-in view of the boxed region exhibited layered distribution of the neurons, whereas the motor neurons are concentrated at the ventral border of the spine. Quantifications of the cell type probability across the spine are shown in the right panel. Scale bar corresponds to 500 µm. (**b**) Results from two MOSTA sections at E16.5 development stage. The mono-layer epidermis (red) structure and pre-epidermal keratinocyte cells (green) under-neath the epidermis layer are highlighted. The scale bar in the insets corresponds to 50 µm.

Next, we computed cell type composition approximated by each cell’s fractional area in the chioso labeling results and compared those results to TOME reference (Fig. 4b, Fig. S3). Note that unlike some of the direct imputation methods^9^, chioso does not enforce an exact population match between the ST data and the reference, and instead rely on the expression pattern alone for inference. This approach allows one to use slightly mismatched references for labeling. For example, in our experiments, TOME was acquired from whole embryo samples, and MOSTA was from thin sections of the embryo samples. Thus, one would not expect the cell composition to be identical between the two datasets. Nevertheless, we found chioso predictions captured the overall trend of composition change over development axis (Fig 4b). For example, in both datasets, we can see rapid declining fractions of chondrocyte and osteoblast progenitors and skeletal muscle progenitors, while simultaneously observe the expansion of connective tissue progenitors and various spinal neuronal types. One notable discrepancy is that chioso reported a significant higher amount of mesenchymal stromal cells than the reference (Fig. 4b). This could reflect the intrinsic bias of the two different technologies, although the exact source of this discrepancy is not immediately clear.

### Comparison with singe-cell segmentation based labeling

To allow for a more quantitative evaluation of the chioso predictions, we performed additional labeling based on single-cell segmentation, using the available DAPI images in MOSTA dataset for segmentation. Specifically, we analyzed an E12.5 section and a E14.5 section. Since the reference dataset contains no E14.5 samples, we used the (slightly mismatched) E13.5 portion of the data for reference. Single-cell masks were computed using a pretrained neural network model^37^ and expression profiles for each segmented cell were extracted. We then performed label transfer from the tome dataset (only the E12.5 and E13.5 subset) using two different software, TACCO^8^ and RCTD^6^. In addition, for chioso results, we also assigned cell-level labels using the same scheme described earlier in the synthetic data section, i.e., by choosing the most frequent pixel-level label within each cell mask. We then compare the results from three different computations (Fig. 6a).

**Figure 6.**
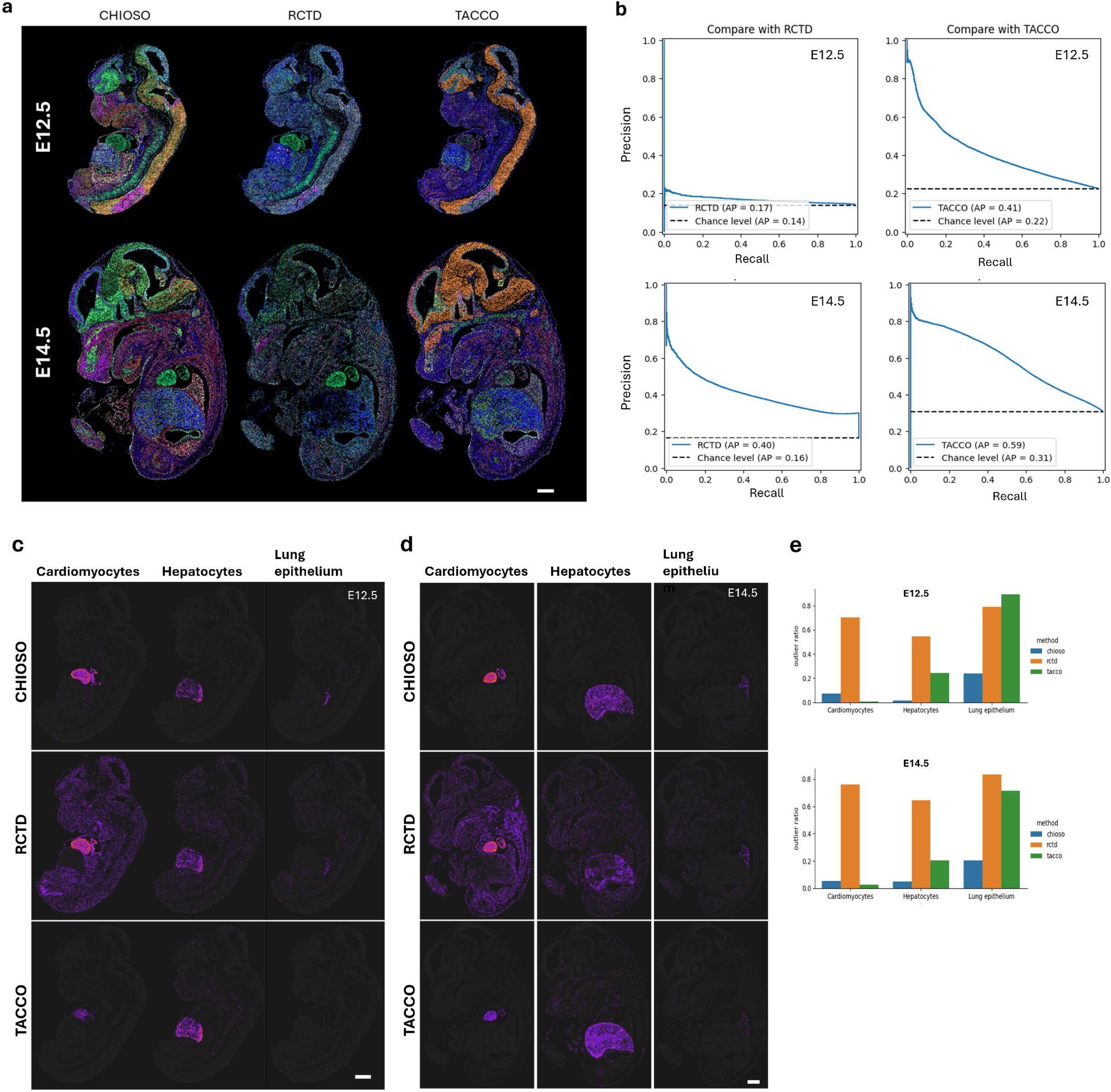
Compare with segmentation-based pipelines. (**a**) Single-cell based labeling results for two MOSTA sections using chioso and two baseline methods. (**b**) Quantification of the consistency between methods via precision-recall curve. TACCO/RCTD results were ranked using their respective score outputs by the original software. (**c)** Comparison of labels of three cell types, cardiomyocytes, hepatocytes, and lung epithelium, that are expected to localize to three specific organs (heart, liver and lung). Results are from the E12.5 section. (**d**) Same as **c** but for the E14.5 section. (**e**) Quantification of results in c & d by quantifying the fraction of “outlier cells” that localize to area outside the specific organs. Scale bar corresponds to 400 µm.

RCTD assign a significant portion (68.8% for E12.5 sample and 64.1% for E14.5 sample) of predictions as “reject” indicating low confidence in the prediction.

Furthermore, agreements between all three results are poor: chioso and TACCO consistency is the highest at 0.22 (i.e., 22% of all cells were given the same label by both methods) for the E12.5 sample and 0.31 for the E14.5 sample; chioso/RCTD consistency is 0.14 for the E12.5 sample and 0.16 for the E14.5 sample; and RCTD/TACCO consistency is 0.13 for both the E12.5 sample and the E14.5 samples. These results reflected the general difficulties of the labeling assay on a noisy ST dataset.

To make an objective comparison among these different results, we specifically examined cell subtypes that are expected to localize to specific organs or anatomical regions and see if the predicted cell locations fell within the expected anatomical regions. Surprisingly, we found chioso predictions consistently have a more clearly defined structure, and better consistency with the expected anatomical knoelwdge of embryos (Fig. 6c & d, Fig. S4). We quantify the results using an outlier detector to classify each cell as either “normal”, representing the cluster of cells within the organ, or “outlier”, representing the disperse cell populations outside the organ region, based on cell locations. In almost all cases, chioso predictions showed the lowest outlier fraction (Fig. 6e). The only exception is for cardiomyocyte, where TACCO predictions showed slightly lower fraction at the cost of detecting a much lower number of total cardiomyocytes and thus having a poor recall. Combined, these results suggested that the segmentation-free approach of chioso can out-perform existing methods that requires single-cell segmentations.

## Discussion

In this paper we present a new algorithm, chioso, designed to annotate large ST dataset at sub-cellular spatial resolution. We believe chioso represents a valuable addition to the current set of toolboxes for ST data analysis, as it has one key desirable feature from the users’ perspective – simple. (1) Since chioso bypasses the single-cell segmentation step completely, it demands a simpler experiment and less experimental time from the user. (2) Chioso is an “end-to-end” algorithm and requires no domain-knowledge dependent data pre-processing (e.g., gene selection) from the user. (3) The chioso output are simply images and therefore compatible with the existing imaging visualization and analysis tools, without requiring specialized software tools. (4) Unlike cell-level labeling, which requires additional computation steps to obtain segmentations of tissue regions, a chioso output is already the tissue segmentations.

We also demonstrated that chioso scale well to large ST dataset with billions of sequencing reads. Note that when the data contains many sections, chioso does not analyze individual sections separately, but instead follow the general machine learning practice of fitting the model globally to all data via stochastic optimization. In such a paradigm, the model training time scales sub-linearly with the dataset size. This allows chioso to scale very efficiently to very large datasets. On the flip side, for very small datasets, chioso may be slower than alternative methods because the fixed training cost, which is potentially a disadvantage of chioso. Another disadvantage of chioso is that it requires specialized computational hardware (GPUs) that may not be (easily) accessible to all biological researchers.

A unique technical feature of chioso is that it is built on the adversarial learning technique. Most reference-based labeling, or “label transfer”, task can be modeled as an optimization problem, which aims to find a mathematical function that convert a target distribution (e.g., distribution of spatial RNA expression profiles) to best match a reference distribution (e.g., distribution of single-cell RNA expression profiles). Currently there are several competing computational frameworks to solve this type of mathematical problems. One framework, that has gained significant popularity^8,38,39^ in recent bioinformatics literature, is the optimum transport (OT) framework. However, for OT to work well, one needs to faithfully represent the *full* distributions of the target and the reference during model fitting. We found it quite difficult to achieve in practice, particularly for data with implied hierarchical structure, such as the ST data. On the other hand, adversarial learning is an alternative strategy to achieve distribution matching. Adversarial learning was originally developed in the field of image synthesis, indicating that it is a method particularly powerful in analyzing spatial data, which is probably why it worked well for our problem too.

## Methods

### Chioso model

Given an ST dataset

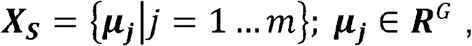

where ***µ***_***j***_ represents the gene expression profile at a specific location, and a labeled reference dataset:

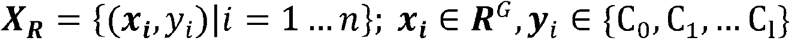

Where ***x***_***i***_ represent the gene expression profile of a specific cell, our aim is to determine the labels *y*_*j*_ ∈ {C_0_, C_1_,⃛} for each ***µ***_***j***_. Here *m* is the number of pixels in the ST dataset, *n* is the number of cells in the reference dataset, *G* is the number of genes, and C_0_, C_1_,⃛ are individual labels, e.g., cell types.

To achieve the labeling, we could seek to find a function 𝔽 that transforms ***X***_***S***_ to a new set that is statistically indistinguishable from {*x*_*i*_}, i.e., the elements of two sets follow the same distribution. For practical reasons, we also want to work in a reduced dimension, since *G* is a very large number (tens of thousands). Hence ℱ has the form:

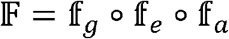

where 𝕗_*e*_ represents a simple linear embedding module for dimension reduction. Although mathematically, it can be described as a simple matrix multiplication,

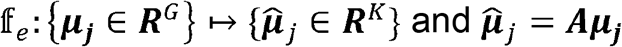

where ***A*** is a real matrix of the shape *G* × *K*, in practice the module is implemented as a lookup table instead, because the input data ***µ***_***j***_ needs to be in a sparse representation to fit into the computer memory. Here, *K* is the embedding dimension much smaller than the number of genes ***G***. Chioso uses a default ***K*** value of 256.

The function 𝕗_*a*_ represent an affine module, whose purpose is to account for the flatform effects, and correct for factors such as the different bias in the library preparations between the ST and the reference dataset.

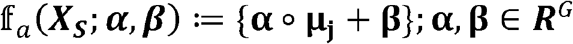

Here º represents element-wise product. Note that chioso does not pre-compute the parameters **α, β** but instead try to learn them together with all other model parameters during model training.

The function 𝕗_*g*_ represents a type of attention mechanism, whose purpose is to learn to integrate nearby pixels to create an expression profile represent the whole cell:

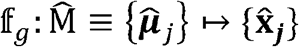

And

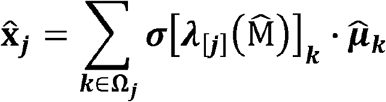

Here **Ω**_***j***_ represents a set of neighboring pixels centered on ***j, σ*(*x*) ≡ *e*^*x*^/ (1+ *e*^*x*^)** is the sigmoid function, and **λ**(·) is a neural network that compute the likelihood that each of those neighboring pixel belongs to the same cell as ***j***. In chioso, we implemented ***λ***(·) using a convolutional neural network (CNN) and use ***λ***[***j***] to denote the CNN output at location ***j***. Note that in practice, we do not need to compute these functions on the whole input dataset simultaneously. Instead, at a time we typically only need to compute on a sample representing a rectangular region of the ST data. In that case, 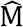 can be
represented as a 3D tensor of the shape *h*×*w*×*C*. We design the ***λ***(·) to output a tensor of *h*×*w*×*C* ′, where *C′* is the cardinality of the set **Ω**_***j***_.

Finally, we also need a machine learning based nonlinear classifier that link cell-level expression profile to a specific cell type label. We implement this as a simple multi-layer perceptron (MLP) neural network parameterized by ***θ***_*p*_ with a softmax activation:

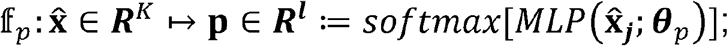

where **p** represents the relative probability for each of the *l* different potential labels.

### Model training

We first determine the values of ***θ***_*p*_, the parameters for 𝕗_*p*_, and ***A***, the parameters for 𝕗_*e*_ via supervised learning using the reference dataset. The objective function is a class-weighted cross-entropy loss function:

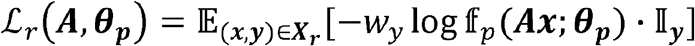

Where *w*_*y*_ is the class weight and 𝕀_*y*_ is the class label in the one-hot vector format.

Next, we determine the rest of the model parameters using an adversarial training scheme. This is achieved by co-training model 𝔽 together with a discriminator 𝔻:***R***^***K***^⟼ [0,1] ⊂***R***, implemented also as an MLP neural network. The training objective is:

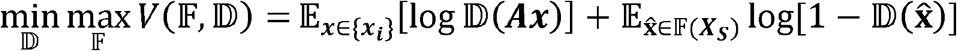

All data presented in the paper were obtained using the default chioso training schedule: By default, the algorithm uses the Adam^40^ optimization algorithm, with L2 weight regularization. The supervised step trains for 10000 batches with a batch size of 128 and a step-wise learning rate scheduler with initial learning rate of 1e-4 and fine-tune learning rate of 1e-5. The adversarial learning used a fixed learning rate of 0.001 for 30000 steps, with options of checkpointing at an earlier training step for downstream inference.

### Evaluation metrics

We primarily use two metrics to evaluate chioso model accuracy. First, to measure labeling accuracy based on cell-level expression, we use balanced accuracy score^41^, defined as:

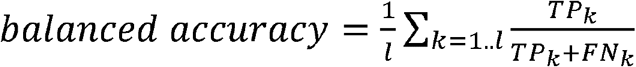

where *l* is the number of classes and *TP* and *FN* denotes the number of true positive and false negative predictions.

To evaluate the ST data labeling accuracy, we used the average precision (AP) metric, defined as:

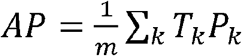

where *m* is the total number of pixels (or cells when evaluating cell-level predictions),*T*_*k*_ is an indicator function of whether the *k*-th prediction, ranked by their confidence scores,
is a true (1) or false (0) one, and *P*_*k*_ is the precision score of the first *k* predictions. AP can also be visually interpreted as the area under the curve of the precision/recall plot and is a value bound between [0, 1].

### Background detection

For visualization purpose, it is typically desirable to display only labels for pixels corresponding to the cell/tissue region and exclude areas where there were no cells. Chioso provides two build-in utilities to achieve this. The first option is an unsupervised method based on thresholding of the RNA counts combined with spatial filtering using a dense CRF filters. To determine the threshold value automatically, we observed that the score output of the model, derived from the discriminator loss values, strongly correlated with cellular locations (e.g. Fig. 2a). Therefore, we implemented a simple optimization algorithm to find the thresholding value by maximizing the intersection-over-union between RNA counts based segmentation and model score images. While this method is easy to use and intuitive, we noticed that in some Steoeoseq data, the background noise is so high that raw RNA counts are in fact higher in non-tissue area. To account for this difficulty, chioso implemented a secondary option based the standard supervised learning framework using CNNs. This option requires a user supplied training dataset, therefore is less convenient to use but potentially can produce better output.

All data visualization in this paper were prepared using the first method, with some manual adjustment of thresholding in individual cases.

### Chioso labeling of synthetic ST data

We use the default chioso configuration, except setting the number of possible cell types to 3, for all model hyperparameters as well as the training schedules.

### Chioso labeling of MOSTA data

We use the default chioso configuration, except setting the number of possible cell types to 68, for all model hyperparameters as well as the training schedules.

### Segmentation based labeling

Single cell segmentation masks are produced on the DAPI images of the original MOSTA dataset for sections E12.5_E1S3 and E14.5_E1S3. We use a deep learning model^37^ trained on the TissueNet dataset^42^ for segmentation. The TissueNet dataset was chosen because it contains data of similar imaging modality (tissue sections stained with DAPI). Only the nuclei segmentation part of the dataset was used. We then computed cell level RNA counts for each section using the single cell masks.

For TACCO labeling, we ran the software using its default configuration. For RCTD labeling, we were unable to run the software for all genes due to out-of-memory error. Instead, we reduced the dataset size by keeping only the top-5000 most highly variable genes using scanpy^43^ software package and performed labeling with the subset of genes using the “doublet” mode.

## Data availability

The original TOME data is available at http://tome.gs.washington.edu/. The original MOSTA data is available at https://db.cngb.org/stomics/mosta/. In addition, we have converted the original data format to hdf5 based custom format, allowing for a much faster on-disk random access suitable for deep-learning pipelines. The converted data files are available as an Hugging Face dataset : https://huggingface.co/datasets/jiyuuchc/mosta. The chioso inference results of MOSTA is available at Mendeley Data with DIO: 10.17632/x99c2gfcwg.1. The synthetic ST dataset generated by this study for method validation is available at Mendeley Data with DOI: 10.17632/6v6smy49r3.1. The segmentation model used to create single-cell mask is available at as a Hugging Face model repository: https://huggingface.co/jiyuuchc/cnsp4-nuc/.

## Code availability

Software implementation of chioso and code to reproduce the results are available at GitHub repository: https://github.com/jiyuuchc/chioso/tree/paper2024.

