## Supplement tables and figures for "Chioso: Segmentation-free Annotation of Spatial Transcriptomics Data at Sub-cellular Resolution via Adversarial Learning": supp_no_fig.docx

**Supplementary Tabel 1**. Capacity and computational complexity of the Chioso model

| Sub-module | Parameters (million) | Million FLOPs per pixel |
| --- | --- | --- |
| Affine | 0.03 | < 0.01 |
| Embed | 7.04 | < 0.01 |
| Predictor | 0.41 | 0.83 |
| Generator | 23.16 | 13.51 |
| Discriminator | 0.26 | 0.54 |
| **Total** | 30.90 | 14.88 |

**Supplementary Tabel 2**. Comparison of different predictor models trained on TOME dataset.

| **Cell Type** | **Chioso Predictor ^[1]^** | | | **MLP w/ PCA Embedding** | | | **Transformer ^[2]^** | | | **support** |
| --- | --- | --- | --- | --- | --- | --- | --- | --- | --- | --- |
|  | precision | recall | f1-score | precision | recall | f1-score | precision | recall | f1-score |  |
| accuracy |  |  | 0.83 |  |  | 0.77 |  |  | 0.83 | 278656 |
| macro | 0.77 | 0.88 | 0.81 | 0.73 | 0.79 | 0.75 | 0.77 | 0.88 | 0.81 | 278656 |
| weighted | 0.84 | 0.83 | 0.83 | 0.78 | 0.77 | 0.77 | 0.84 | 0.83 | 0.83 | 278656 |

[1] K=256

[2] Six-layer transformer model with 2048 tokens of 256 dimensions.

**Supplemental Figure Captions**

**Figure S1**. Predictor performance versus model embed dimensions ***K***. The model training achieves lower loss (a) and higher validation balanced accuracy (b) when the *K* value was increased from 128 to 256. Further increase to 512 resulted in lower loss but not in validation accuracies. All tests were done using the TOME dataset.

**Figure S2.** Additional visualization of MOSTA results. (**a**) Mouse embryo at E16.5 development stage. (**b**) Mouse embryo at E13.5 development stage. **(c)** mouse embryo at E11.5 development stage.

**Figure S3.** Complex spatial organization of spinal cord. Chioso labeling of a E16.5 sample (top) and E13.5 sample (bottom) revealed the spatial organizations of three dominant cell types (excitatory neuron, inhibitory neuron and motor neuron) within the region. Zoomed-in view of the boxed region exhibited layered distribution of the neurons, whereas the motor neurons are concentrated at the ventral border of the spine. Quantifications of the cell type probability across the spine are shown in the right panel. Scale bar corresponds to 500 μm.

**Figure S4.** Chioso exhibit high spatial resolution in labeling. Shown here are results from two MOSTA sections at E16.5 (left) and E14.5 (right) development stage respectively. The mono-layer epidermis (red) structure and pre-epidermal keratinocyte cells (green) under-neath the epidermis layer are highlighted. The scale bar corresponds to 50 μm.

**Figure S5.** Chioso prediction of MOSTA cell compositions over the development. (a) Average number of pixels (2 μm) for each cell type showing embryo growth over time. (b) Normalized cell compositions approximated by area fraction.

**Figure S6**. Additional comparison between chioso and two baseline methods (RCTD and TACCO) that requires single-cell segmentation. The results for the E12.5 section is on top in (**a**) and results for the E14.5 section is at the bottom (b). Eight cell types were chosen based on their recognizable anatomical features. The first three corresponds to specific organs. The next four corresponds to specific regions of nervous system. The last one represents the skeletal structure.
