## Supplementary figures and images for "Chioso: Segmentation-free Annotation of Spatial Transcriptomics Data at Sub-cellular Resolution via Adversarial Learning"

### FigS1.png

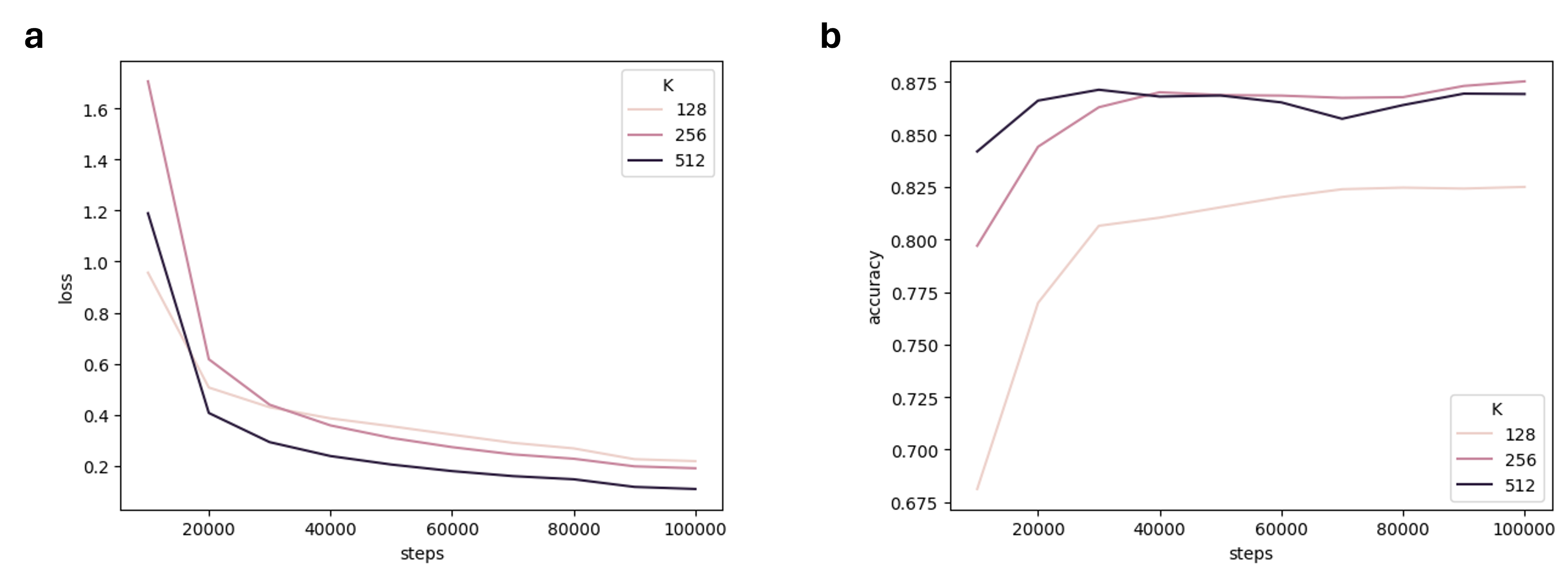

### FigS2a.png

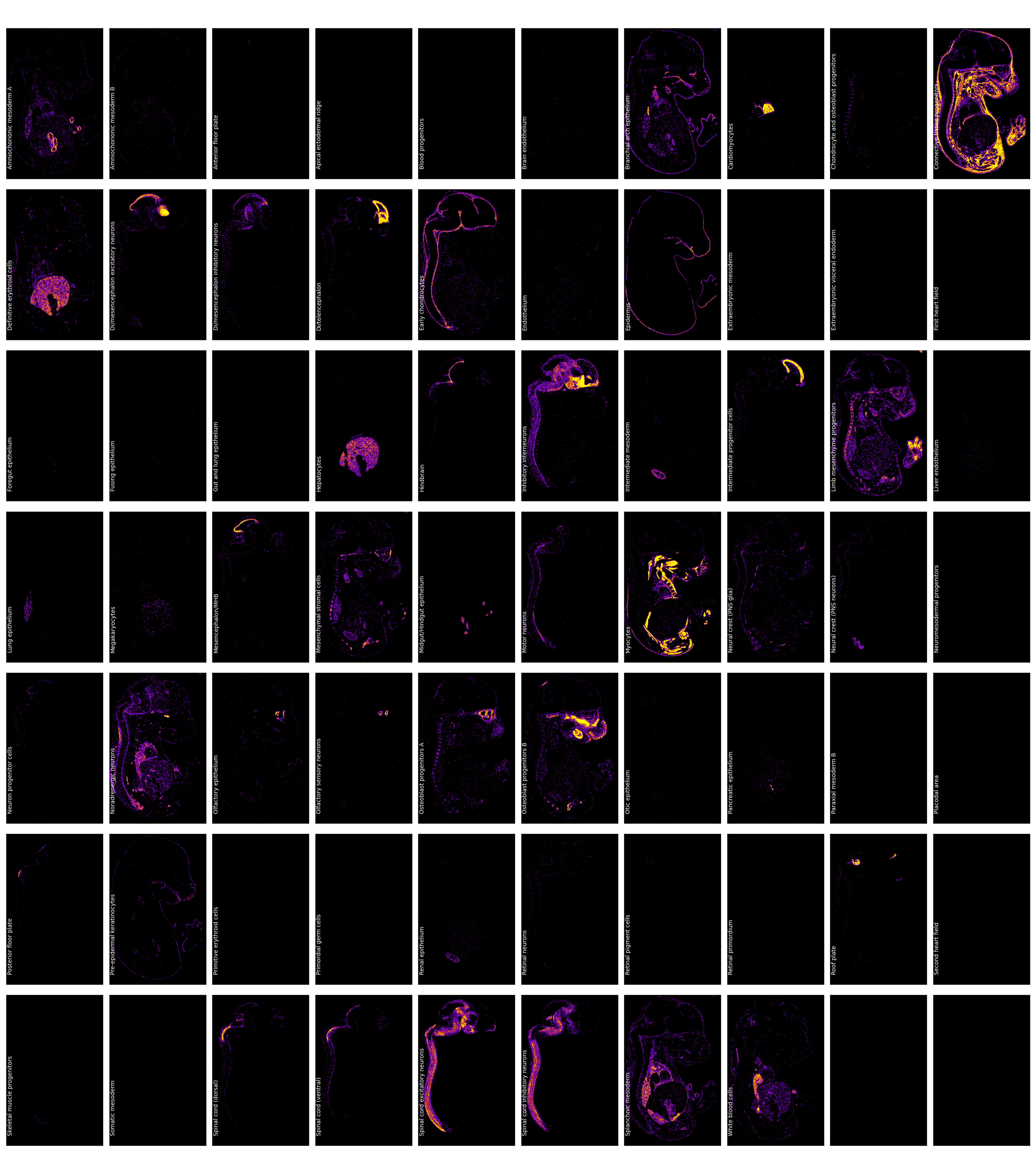

### FigS2b.png

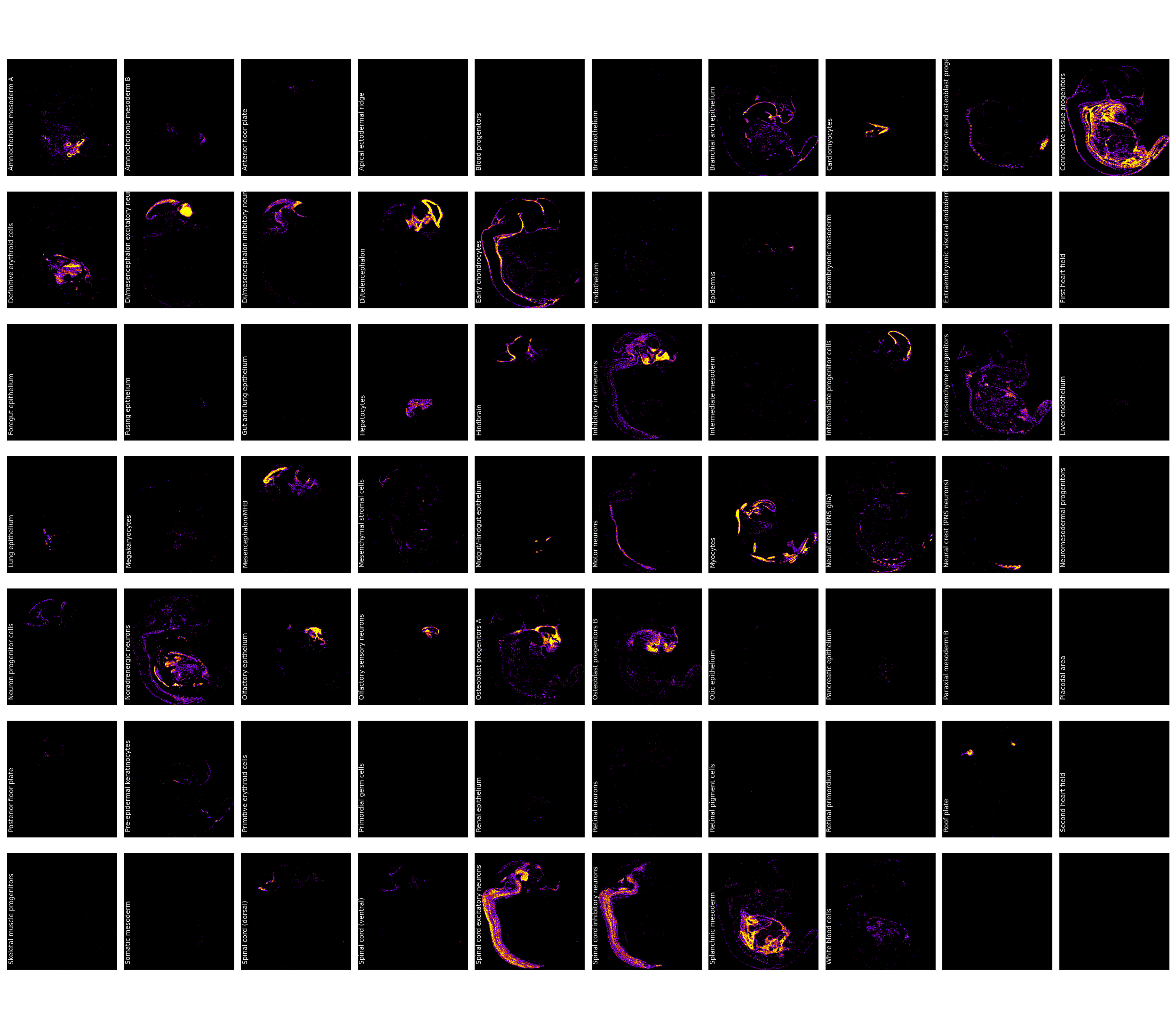

### FigS2c.png

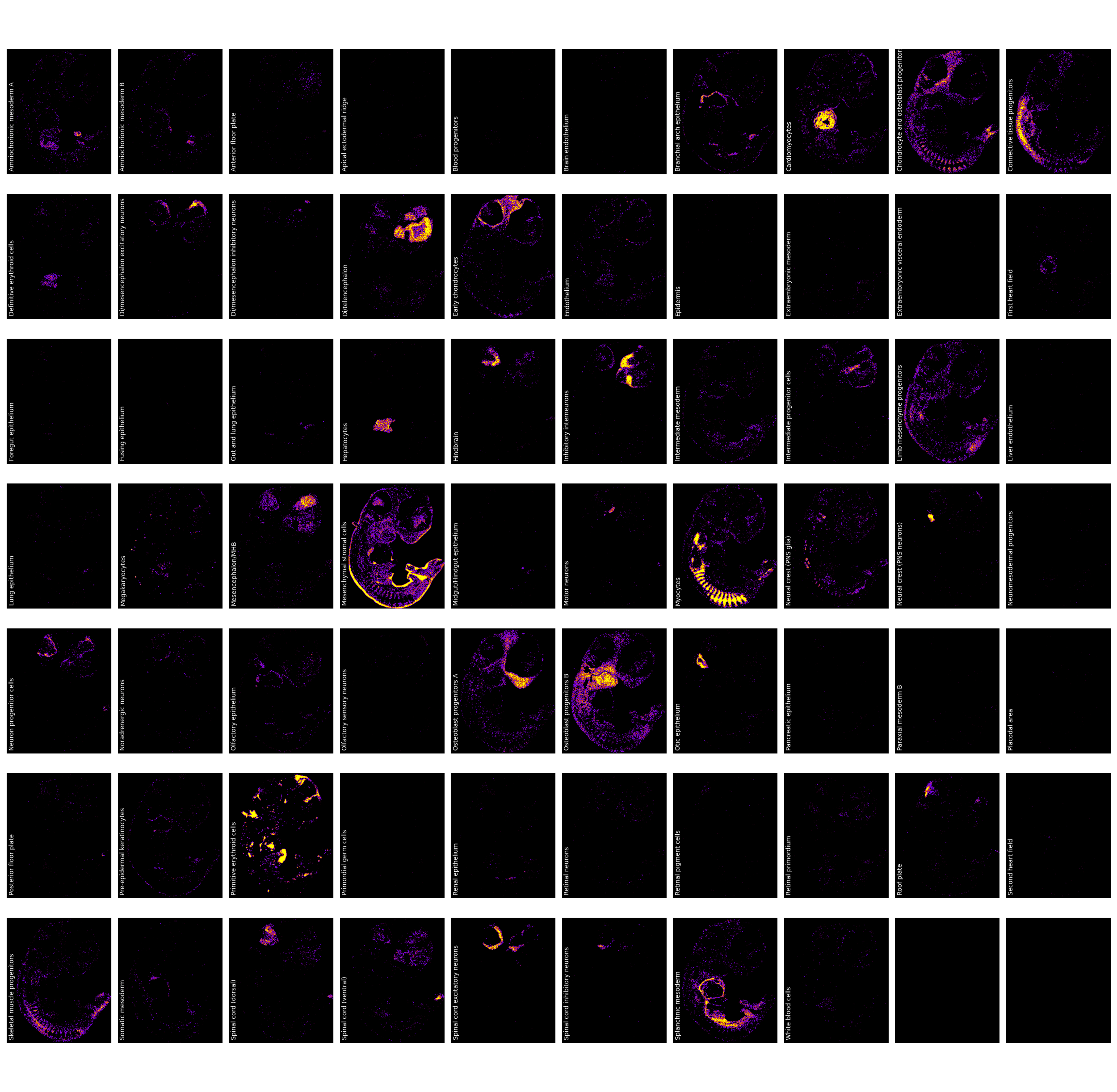

### FigS3.png

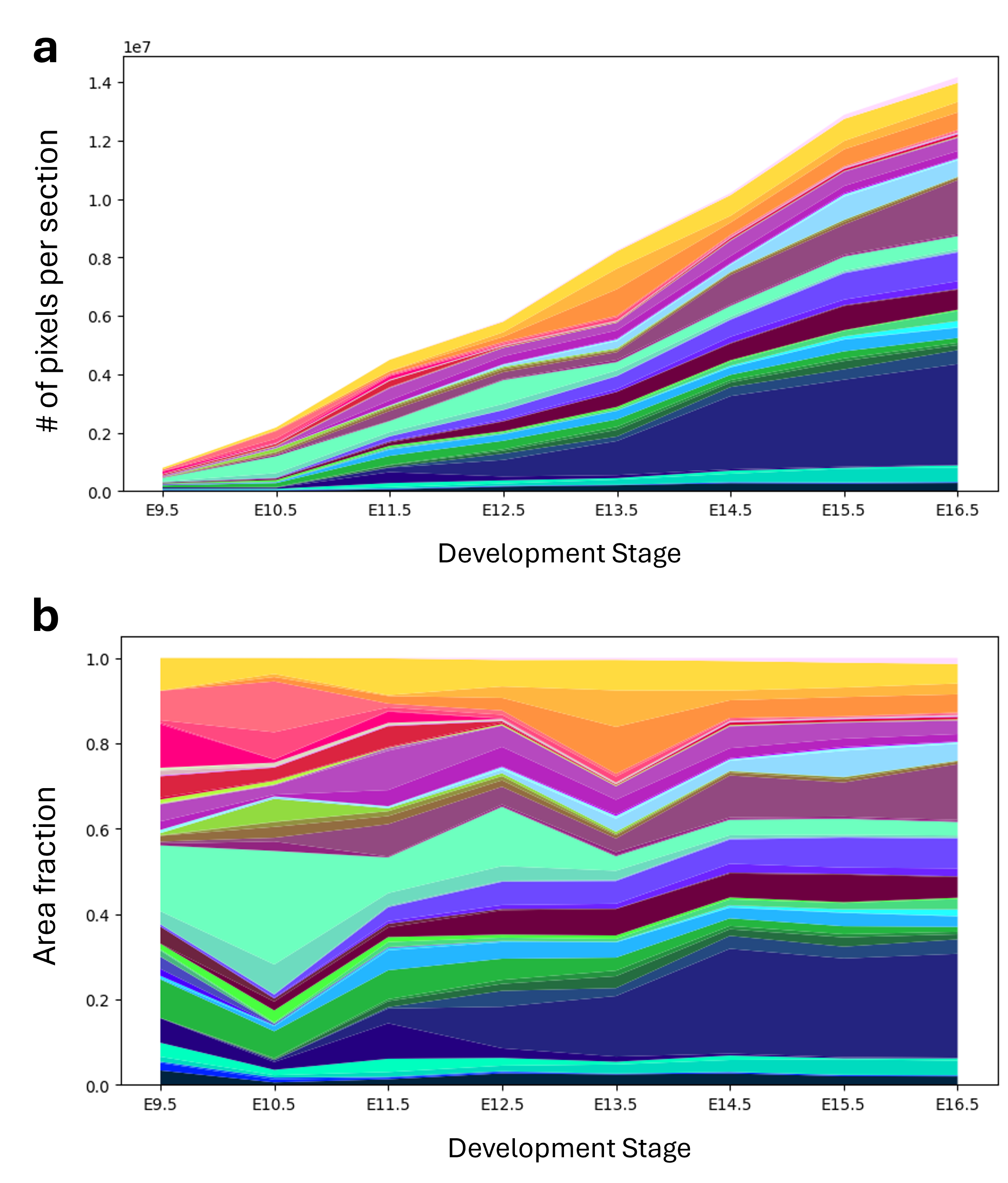

### FigS4.png

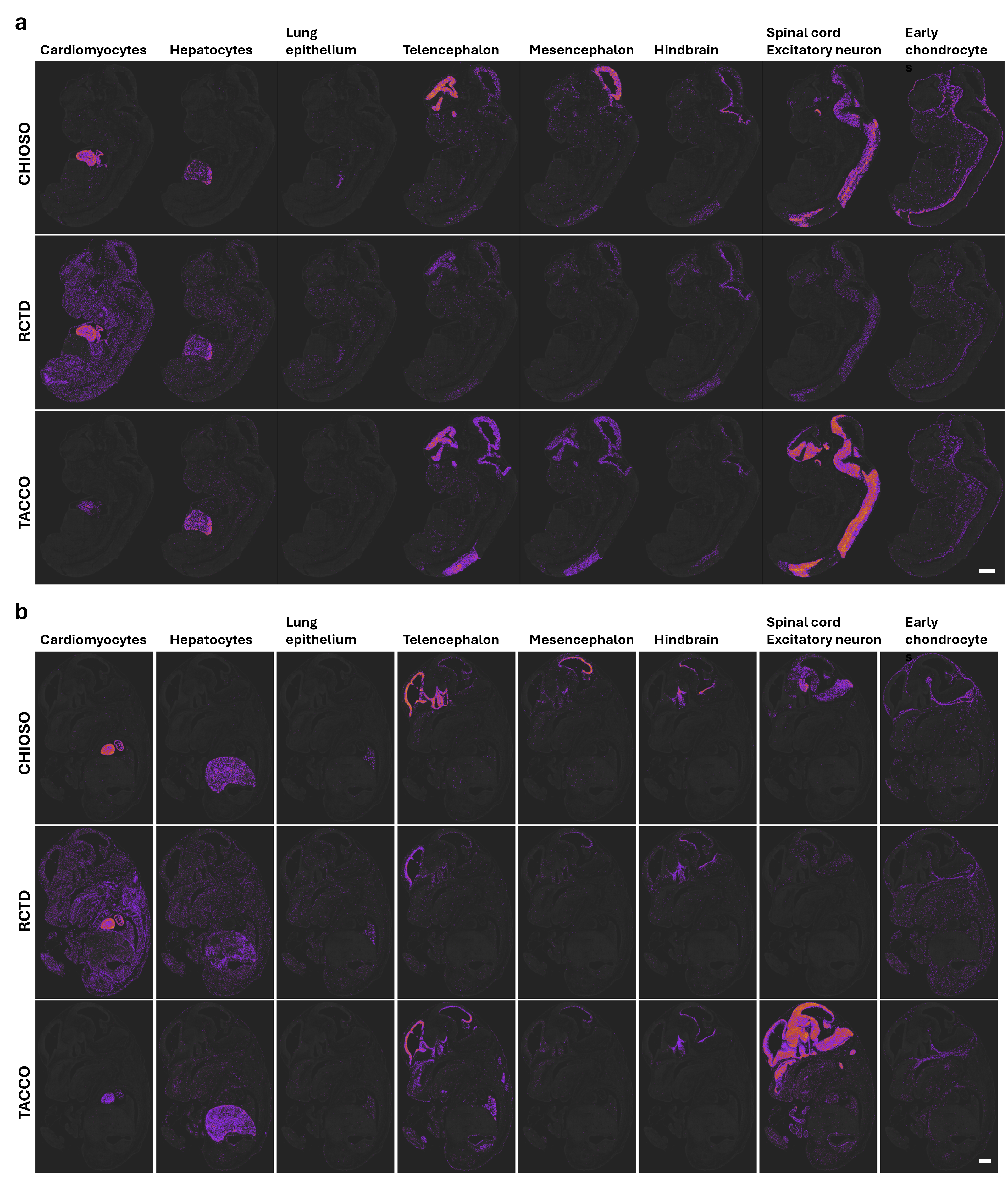
